# Quantitative Visualization of Hypoxia and Proliferation Gradients Within Histological Tissue Sections

**DOI:** 10.1101/807875

**Authors:** Mark Zaidi, Fred Fu, Dan Cojocari, Trevor D. McKee, Bradly G. Wouters

**Affiliations:** Department of Radiation Oncology and Medical Biophysics, University of Toronto, Toronto, ON, Canada; STTARR Innovation Centre, Princess Margaret Cancer Centre, University Health Network, Toronto, ON, Canada

**Keywords:** Hypoxia, tumor microenvironment, digital pathology, immunofluorescence, image analysis, distance mapping, biological gradient, tissue cytometry

## Abstract

The formation of hypoxic microenvironments within solid tumors is known to contribute to radiation resistance, chemotherapy resistance, immune suppression, increased metastasis, and an overall poor prognosis. It is therefore crucial to understand the spatial and molecular mechanisms that contribute to tumor hypoxia formation to improve the efficacy of radiation treatment, develop hypoxia-directed therapies, and increase patient survival. The objective of this study is to present a number of complementary novel methods for quantifying tumor hypoxia and proliferation, especially in relation to the location of perfused blood vessels.

Multiplexed immunofluorescence staining can produce whole slide scanned image datasets that are amenable for computational pathology analysis. A standard marker analysis strategy is to take a positive pixel count approach, in which a threshold for positive stain is used to compute a positive area fraction for hypoxia. This work is a reassessment of that approach, utilizing not only cell segmentation but also distance to nearest blood vessel in order to incorporate spatial information into the analysis. We describe a reproducible pipeline for the visualization and quantitative analysis of hypoxia using a vessel distance analysis approach. This methodological pipeline can serve to further elucidate the relationship between vessel distance and microenvironment-linked markers such as hypoxia and proliferation, can help to quantify parameters relating to oxygen consumption and hypoxic tolerance in tissues, as well as potentially serve as a hypothesis generating tool for future studies testing hypoxia-linked markers.

## Introduction

Solid tumors are often characterized by heterogeneity in key microenvironmental features, including variations in cell type (tumor, stromal or immune cell content), availability of nutrients, and oxygenation. Micro-regional changes in oxygenation are due to mismatches in metabolic consumption relative to oxygen supply to tumor cells. Typically, areas of hypoxia are defined as those below a threshold of oxygen required to confer a specific biological or therapeutic impact. The presence of regions with low oxygen partial pressure (pO_2_) of ≤ 20 mmHg confers a greater resistance to radiation therapy *(Alper and Bryant, 1974)* and conventional chemotherapeutics *(Gomez, 2016)*, and are correlated with lower patient survival *(Rudat et al., 2001)*. The vasculature of solid tumors is often abnormal due to either tortuous vasculature formation or vascular collapse *(Milosevic et al., 1999)*. Regions near blood vessel capillaries can be normoxic, with cells experiencing hypoxia as pO_2_ decreases due to oxygen metabolism away from vessels. This eventually leads to necrosis when pO_2_ becomes insufficient to support tumor cell viability.

The physiological relationship between blood vessel distance, presence of viable tumor tissue, and metabolic activity was first described in *(Thomlinson and Gray, 1955)* which showed that the shape of the oxygen gradient is determined by the metabolic oxygen demand within tissue. Although rates of oxygen consumption can vary substantially depending on tissue type, oxygen gradients on the order of 150 μm from supply to anoxia are typically reported *(Haugland et al., 2002)*. This phenomenon is commonly referred to as chronic hypoxia because the match between oxygen diffusion and consumption is relatively stable and thus the exposure of cells to hypoxia is long lived *(Bayer and Vaupel, 2012)*. Tumor cells can also become acutely hypoxic due to transient changes or occlusion of blood vessels. It is important to distinguish chronic and acute hypoxia because the length and severity of hypoxia (and reoxygenation) can have different implications for treatment efficacy and hence the impact of hypoxia on patient prognosis *(Vaupel and Meyer, 2007)*.

Coupled with contrast agents and tracers, hypoxia can be imaged in a variety of imaging modalities such as PET, MRS, MRI, NIRS, and EPR. While all share the advantage of being non-invasive, they lack sufficient spatial resolution to accurately detect patterns of hypoxia within the tumor microenvironment *(Jensen, 2009; Carreau et al., 2011)*. In a clinical setting, microelectrodes such as the Eppendorf oxygen probe can measure extracellular pO_2_ and pH (*Rudat et al., 2001, Milosevic et al., 2012)*. While providing a quantitative readout of oxygen concentrations, they are technically demanding to use and also impart poor spatial resolution. Histological staining against exogenously administered markers of hypoxia provide much higher spatial resolution necessary to image hypoxia micro-regional distribution (*Mirabello et al., 2018*). A common method to quantify hypoxia in stained histological sections is to apply binary thresholding. The fraction of pixels above a predefined threshold measures hypoxia-positive area as a percentage of tumor area *(Loukas et al., 2004)*. While other cellular markers can be quantified this way, hypoxia should not be because it is a gradient rather than a binary metric *(Russell et al., 2009)*. Since hypoxia is a continuous gradient within tissues, there is no universally accepted threshold for discriminating hypoxia from normoxia. It is not uncommon for studies to use arbitrarily determined threshold values *(Urtasun et al., 1986)*.

Several studies have employed different methods for measuring hypoxia gradients relative to blood vessels. One popular method is vessel distance analysis (VDA), particularly given development of image analysis platforms capable of this type of analysis. VDA computes mean marker intensity in an image object (pixel, segmented cell, or some other region of interest (ROI)), as a function of distance to a vessel. In *(Rijken et al, 2000)*, VDA was performed in a human glioma xenograft using two different hypoxia markers (NITP and pimonidazole). Concentric rings were generated around perfused blood vessels to calculate mean intensity of the two hypoxia markers in each ring. This study used 50 μm-wide concentric rings as the distance bin, reporting broad ranges of maximal hypoxia; the use of smaller (~10 μm) distance bins would have more accurately pinpointed the distance at which maximal hypoxia staining would occur. A similar analysis was performed on human HNSCC *(Wiffels et al., 2000)* using PAL-E vimentin as a vasculature marker and pimonidazole as the hypoxia marker, with a serial hematoxylin and eosin (H&E) stained section used to delineate tumor area and exclude necrotic regions. Although one of the goals of this study was to measure proliferation, they did so indirectly by measuring hypoxia and inferring the normoxic regions to be proliferative. A more accurate measure of proliferation would be to quantify staining for either an exogenous (EdU or BrdU) or endogenous (Ki67) marker. *Swinson et al.* stained for CA9, an endogenous marker for hypoxia, and manually measured distribution of CA9-positive cells relative to CD34-defined vessels *(Swinson et al., 2003)*. In this study, the authors excluded any vessel that had been cut on its longitudinal axes, as it was easier to measure oxygen gradients from perpendicular vessels. However, removing potential sources of oxygenation from analysis could lead to a misinterpretation of the tumor being less oxygen dependent. *Primeau et al.* used VDA to measure the chemotherapeutic doxorubicin relative to all CD31-positive vessels, however perfusion was not assessed *(Primeau et al., 2005)*. In addition to hypoxia, cellular proliferation has also been shown to be oxygen dependent *(Tannock, 1968). Russell et al.* co-injected two hypoxia markers with perfusion marker and measured hypoxia using cumulative histograms of marker area at different positive staining thresholds *(Russell et al., 2009)*. However, analysis was performed using positive pixel fractions and spatial localization was not studied (*Russell et al., 2009)*. By using a perfusion marker to identify functional vessels, a better understanding of diffusion-limited hypoxia can be achieved.

Here we compare a number of data visualization and analysis methodologies for approaching the quantitative analysis of tissue hypoxia at a cellular level. We describe image ROI and cell segmentation, generation of distance maps to intra-tumor spatial features, scatterplots to interpret marker intensity correlations, and distance bin histograms to interrogate hypoxia distance relationships to tissue features like perfused vessels and necrosis. We find that the extent of hypoxia can be affected by the diffusion and consumption of oxygen within tissues, as well as the tolerance of cells towards surviving in low oxygen conditions. A combination of measurement of the gradient of hypoxia versus distance to vessel, or distance to necrosis, as well as per-cell scatterplots that relate markers of interest (e.g. proliferation and hypoxia), can provide a more robust quantification of hypoxia within tissue sections.

## Methods

### Immunofluorescence Histology

NOD scid gamma mice bearing a KP4 pancreatic cancer cell line xenograft were administered intraperitoneal injection of 400 μl of 2.5 mg/ml EdU (proliferation marker) and 250 μl of 10 mg/ml EF5 (hypoxia marker), 30 min and 3 hours, respectively, prior to tumor excision. In addition to KP4 cell line, we have found that other pancreatic (PANC1 and B×PC3) and colorectal (HCT116 and UM-SCC-74B) cell lines are suitable for vessel distance analysis (Cojocari, 2017), as are colorectal patient derived xenografts (Haynes et al, 2018). To determine which blood vessels were actively perfused at the time of tumor excision, 100 μl of 10 mg/ml Hoechst 33342 was injected into the tail vein of the mouse 1 minute prior to tumor excision. Tumors were embedded in optimal cutting temperature compound (OCT), snap frozen, and sectioned using a cryomicrotome at 5 μm thickness. Unstained sections were scanned for Hoechst using a TissueScope 4000 (Huron Technologies) at 10x magnification with a DAPI filter. Cy3-conjugated anti-EF5 (1/120 dilution of 1μg/1ml stock) and Cy3-conjugated Click-IT EdU reagents (Invitrogen, cat. C10634) were utilized to label hypoxia and proliferation, respectively. Platelet endothelial cell adhesion molecule (CD31), expressed on the surface of blood vessels, was stained for using a rat anti-mouse CD31 antibody (1/200 dilution, BD PharMigen PECAM-13.3 cat. 553370 lot 86580). Secondary AF488-conjugated goat anti-rat (Invitrogen, cat. A11006) was used against the rat anti-mouse CD31 antibody. DAPI nuclear counterstain was applied at 1 μg/ml concentration. Slides were scanned for EdU, EF5, and DAPI using Cy5, Cy3, and DAPI filters, and then rescanned for EF5, CD31, and DAPI using Cy3, FITC, and DAPI filters, respectively. Slides were subsequently stained with hematoxylin and eosin (H&E) to differentiate different tissue regions based on morphology. Brightfield scans were taken with an Aperio AT2 whole slide scanner at 20× magnification. In the end, four separate images were obtained: a single-channel Hoechst image, a EdU-EF5-DAPI immunofluorescence image, a EF5-CD31-DAPI immunofluorescence image, and a brightfield H&E image. The entire process of staining and image acquisition is shown in **Supplementary Figure 1**.

### Image Analysis Methodology

The EF5-CD31-DAPI and EdU-CD31-DAPI RGB images were converted into single-channel grayscale TIFF images. The H&E image was separated into red, green, and blue grayscale TIFF images, and intensity was inverted to produce dark backgrounds, for intensity-based alignment. Semi-automated intensity-based image registration was performed using a similarity transform, which allowed for translation, rotation, and scaling, but not shearing of the images. The intensity-inverted H&E image was used as the target static image for registration. Alignment was manually inspected, and manual control-point alignment was performed if intensity-based alignment was poor. Aligned images were exported as a series of uncompressed 8-bit single-channel TIFF images which were subsequently imported into Definiens Tissue Studio (Definiens Inc, Munich Germany) for image segmentation and classification. Similar workflows could be achieved in several other digital pathology platforms.

Tissue was separated from background using the H&E image as reference. Image subsets were used to train the machine learning classifier to identify regions of interest (ROIs) for hypoxia, necrosis, viable tumor, empty space, and perfusion. This was done using EF5, DAPI, and Hoechst image layers as input. Within 200×200 μm sample subsets, we manually annotated samples of each ROI to train the proprietary Definiens classifier. Hypoxia, perfusion and necrosis were annotated based on high EF5 intensity, high Hoechst intensity, and regions of increased eosin staining and condensed nuclei in the H&E image, respectively; and all remaining non-artifactual (i.e. excluding stroma, musculature, and folds) tissue regions were defined as viable tumor. After a reasonable classification was achieved on the training data set, the trained classifier was applied across all images. Manual correction was used to correct misclassified regions and remove artifacts such as folded tissue present on the slide. ROI annotations were reviewed by a trained histopathologist.

To perform vessel distance analysis, cells were first segmented by detecting nuclei on the DAPI channel, which was performed in the hypoxia, perfusion, and viable tumor ROIs. A watershed algorithm disconnected closely-packed nuclei and a size threshold was applied to exclude nuclei less than 23 μm^2^ in area. Vessel detection was performed on CD31 channel, identifying vessels greater than 5 μm^2^ in area. Following batch processing, the resulting cell and vessel image objects were imported into Definiens Developer XD. A distance map calculating distance to the center of each vessel produced a grayscale image layer with intensity increasing proportionally away from CD31 positive vessels. Distance maps to Hoechst and necrosis ROIs were also generated. For each image, a table of image objects (cells) was exported, along with per-cell features including centroid coordinates; nucleus and cell area; mean marker intensity for CD31, DAPI, EF5, EdU, and Hoechst; and distances to all vessels, perfused (Hoechst) regions, and necrosis. Data visualization and distance bin generation was performed in MATLAB.

ROI-based distance analysis was achieved by defining concentric rings with a width of 10 μm around the Hoechst ROI extending outward. The relative fraction of hypoxia, viable tumor, and necrotic ROIs were calculated in each ring up to a distance of 700 μm. Marker Area Detection (MAD) and Cellular Classification (CC) were employed to detect positive staining. In MAD, individual pixels were grouped into negative, low, medium, and high categories based on EF5 intensity using user-defined thresholds. Thresholds were selected by the operator such that the negative-low threshold would be a first-pass threshold to mark any cells with no observable EF5 staining as negative. The threshold separating low from medium was set higher to identify cells of intermediate EF5 staining, with that separating medium from high used to identify cells with the most intense EF5 staining. Connected regions less than 10 μm^2^ were excluded from analysis. CC was performed by dilating previously detected nuclei by 5 μm to simulate the area of a cell in the absence of a membrane marker, with user-defined thresholds applied to mean EF5 intensity inside the cell. Please refer to the data availability statement of this paper for the code used and further details on the methodology.

## Results

### Comparison of Image Segmentation Methods for Thresholding Hypoxia

Our first approach to assess the amount of hypoxia present was to apply a series of three intensity thresholds to the viable tissue area using both the MAD and CC methods. In CC, by segmenting cells using nuclear stain and morphology, extracellular staining can be filtered out. Cell simulation around each nucleus captures cytoplasmic staining. **Figure 1** shows a comparison between MAD (**Figure 1B**) and CC (**Figure 1C**). While MAD regions seem to be relatively homogeneously distributed in hypoxic regions, CC regions appear to emerge as a concentric gradient of low to high EF5 staining from perfused regions stained with Hoechst. Necrotic regions were excluded from analysis. We utilized the same thresholds in MAD and CC analysis, **Figure 1** shows roughly a threefold increase in percent positivity using CC compared to MAD.

**Figure 1.**
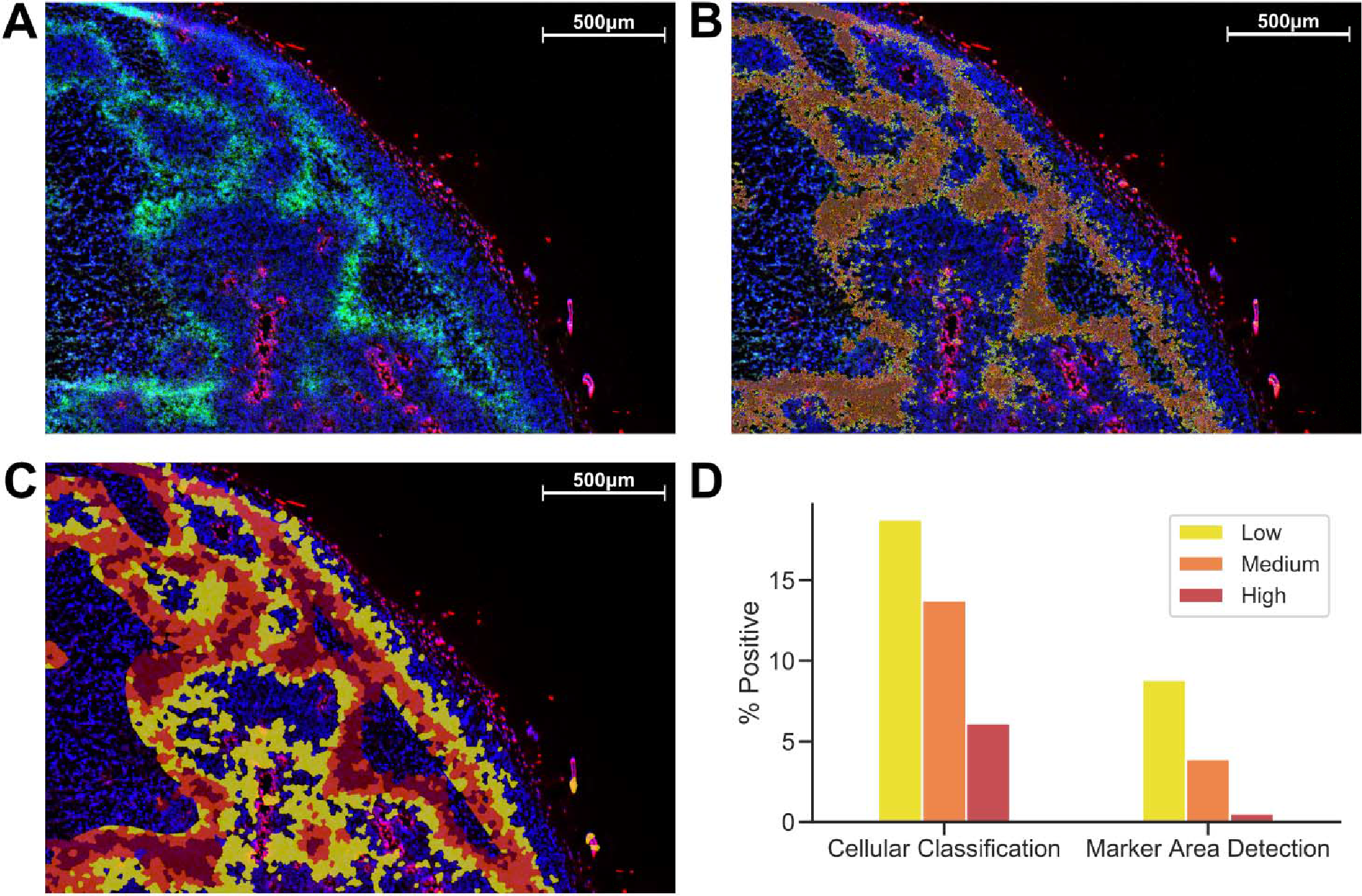
Comparison of Cellular Classification and Marker Area Detection. **(A)** Immunofluorescent staining with DAPI in blue, Hoechst in red and EF5 in green. **(B)** MAD with low (yellow), medium (orange) and high (red) hypoxic areas. **(C)** CC with the same overlay classification as previously used for MAD. **(D)** Comparison of percent positive scores for each EF5 intensity grouping in CC compared to MAD.

### Region of Interest-based Distance Analysis

Another strategy is to define ROIs that comprise the tissue area and measure distances between distinct regions. We incorporate ROIs into our analysis methodology by excluding regions of necrosis from viable tumor during the preparation of our images for processing. In **Figure 2B**, Hoechst perfusion was utilized to identify regions of perfusion surrounding blood vessels and a hypoxia ROI was identified by the presence of EF5 staining. The distance to the nearest Hoechst positive region was calculated as a separate image layer (**Figure 2C**). **Figure 2D** reports the tissue composition for viable tumor, hypoxia and necrosis ROIs as a function of distance from Hoechst perfusion.

**Figure 2.**
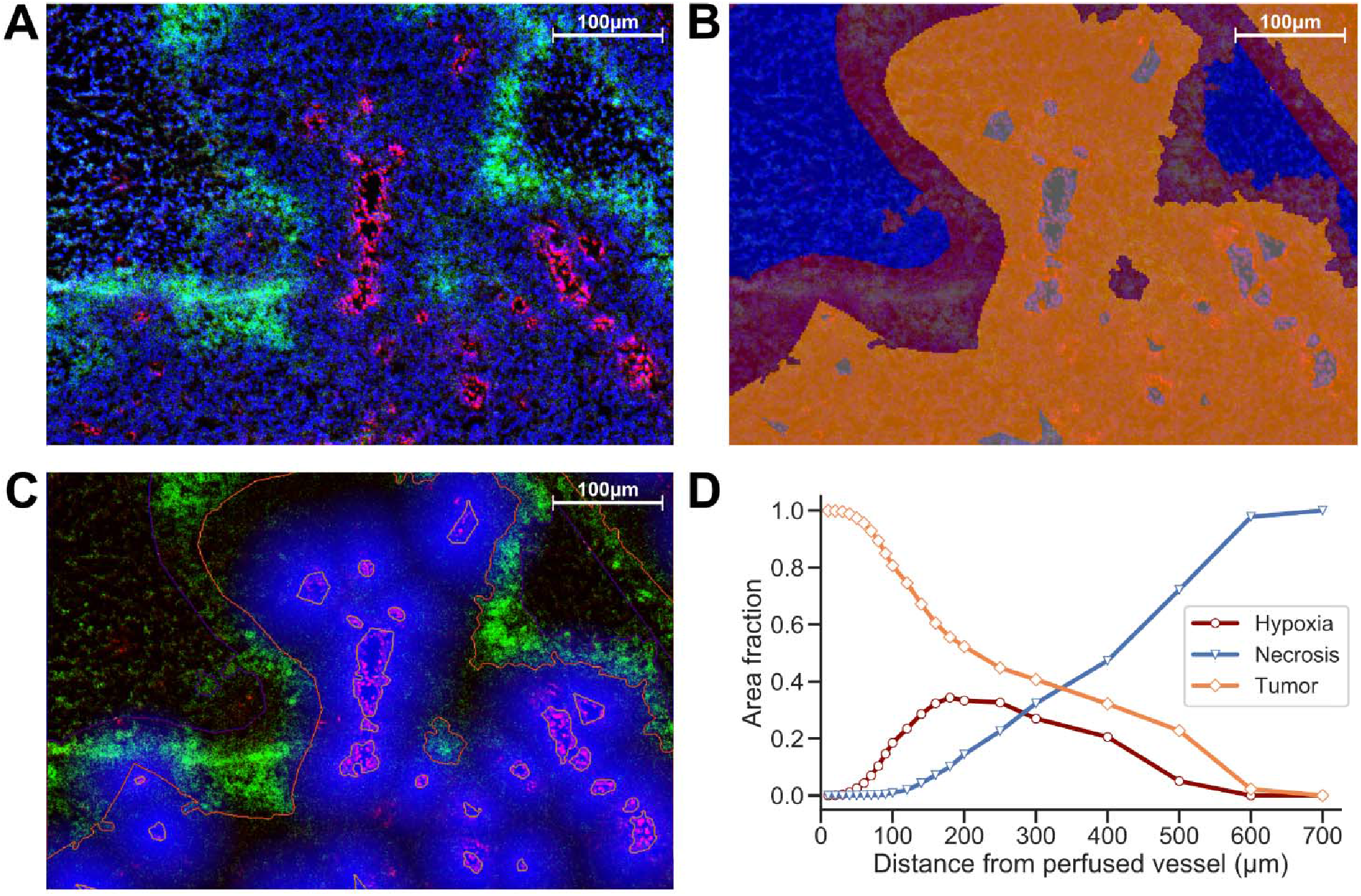
(A) Immunofluorescence image of KP4 xenograft. Stains include CD31 (blue), EdU (red), and EF5 (green). **(B)** ROI overlay of necrosis (blue), hypoxia (red), viable tumor (orange), and perfusion (gray). **(C)** Distance map to Hoechst perfusion area is shown in blue, with intensity decreasing in proportion to the distance away from the Hoechst positive region. **(D)** Viable tumor area decreasing and necrotic area increasing as a function of distance from perfused vessels, with hypoxic area peaking near 180 μm.

### Marker Intensity Histograms & Density Scatter Plot Visualization

To visualize colocalization between multiple markers, dual-marker density scatterplots (similar to those commonly used in flow cytometry analysis) were produced with each axis corresponding to an individual cell’s EF5 and EdU intensity. **Figure 3A** shows a typical flow cytometry-like density scatterplot of per-cell EF5 and EdU intensities, with colors indicating cell density and gates drawn at mean plus one standard deviation for each marker. In **Figure 3B,** transparency of each point is set proportional to the total number of cells. Regions with higher density will appear more opaque than lower density regions. We then incorporate spatial information into this visualization by coloring points according to their distance to the nearest perfused vessel. It can be seen that the subpopulation of EF5-positive cells have a greater average distance. However, it is difficult to observe a distance relationship in the EdU-positive subpopulation.

**Figure 3.**
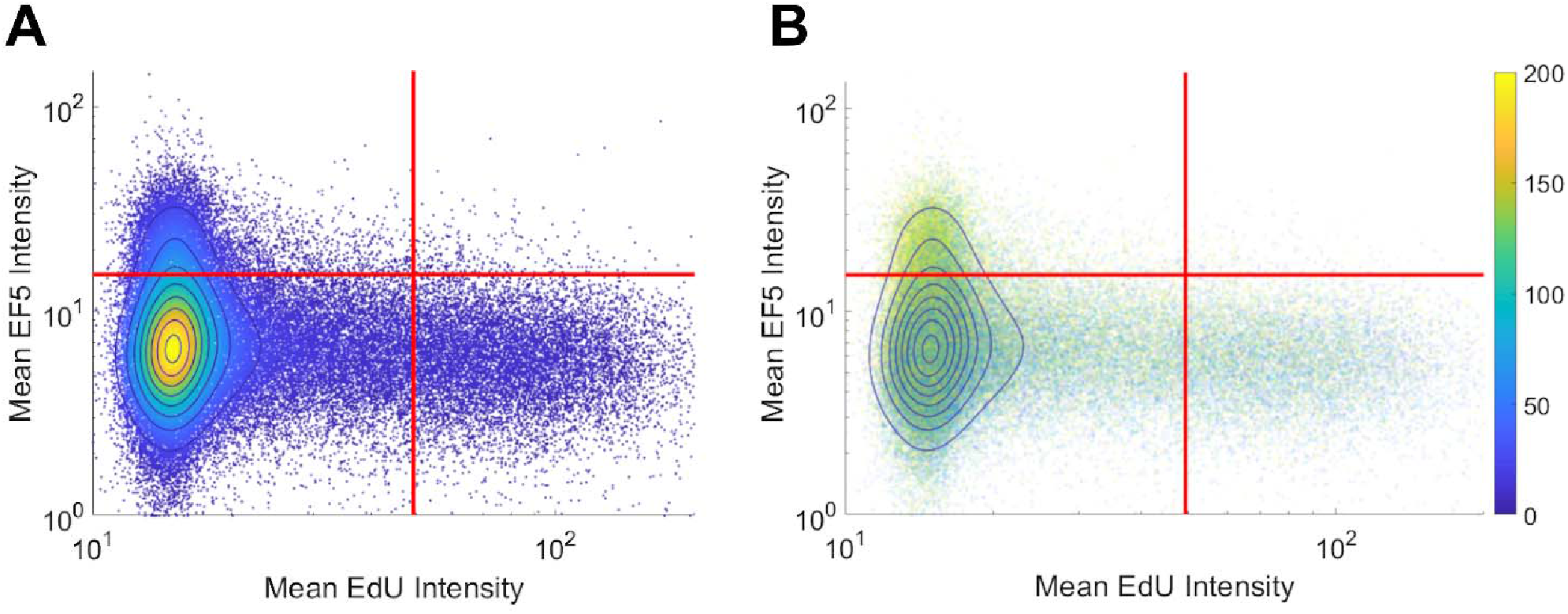
Per-cell scatterplot of hypoxia marker (EF5, vertical axis) versus proliferation marker (EdU, horizontal axis). Regions of increasing density are shown with contour lines in both images. **(A)** density is also represented in color, while in **(B)** individual cells are colored by their individual distances to the blood vessel regions (distances, in microns, shown on colorbar).

### Hypoxia and Proliferation Gradients Relative to Vessels and Necrosis

Distance maps were generated from perfusion (Hoechst) ROI to identify per-cell distances to the nearest perfused vessel. Distance to all CD31-positive vessels, and distance to necrotic ROIs, were also generated on a per-cell basis. To validate the accuracy of cell and vessel segmentation, manual counts were performed in random 350×350 μm tiles and compared to Definiens algorithm-generated counts. The percent errors for cell and vessel detection were found to be 4.7% and 5.3%, respectively, indicating that reliable segmentation was achieved. From this data, cells were binned into uniform concentric distance regions. **Figure 4** presents cell count histograms per bin, which aid in interpretation of the distribution of mean intensity and the percent positive cell intensity for the EdU proliferation marker, and the mean intensity of the EF5 hypoxia marker, versus distance.

**Figure 4.**
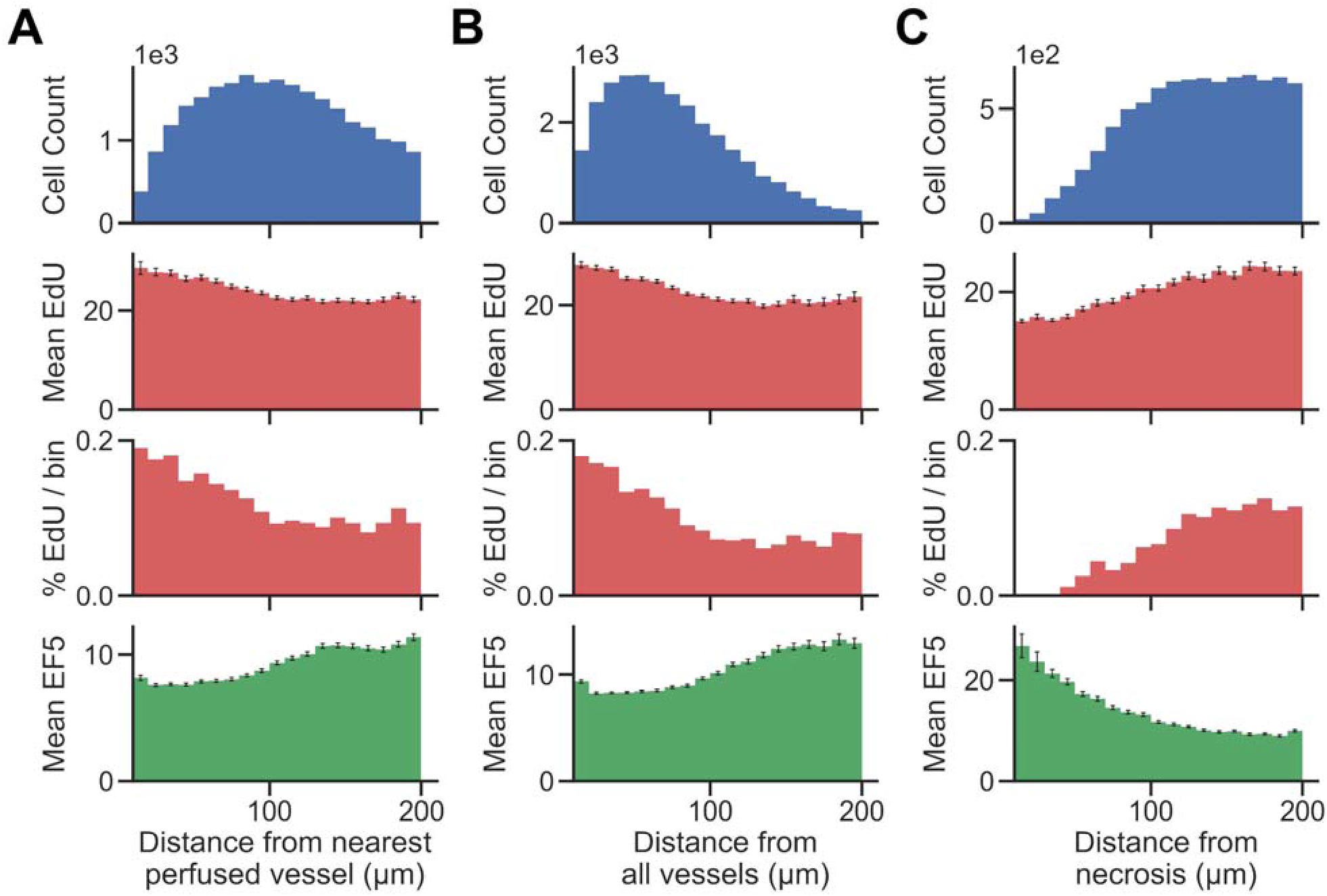
Histograms showing the total number of cells (top) present within uniform distance bins measured from the region of interest (**(A)** distance from the nearest perfused blood vessel, **(B)** distance from all blood vessels, and **(C)** distance from necrosis) in the image. The mean EdU intensity within all of the cells identified in these regions is shown (second from top, measured in arbitrary fluorescence staining units), as well as the mean EF5 staining intensity (bottom). Error bars correspond to the standard error of the mean of the marker intensity in each distance bin. We also utilize a threshold for positivity for EdU signal to identify EdU positive cells, and display the percent of cells in each distance bin that is positive for the EdU marker (second from bottom).

## Discussion

The simplest quantification method for immunostained markers on tissue sections is to apply a pixel-level threshold on the stain of interest. While less algorithmically complex than cell segmentation, MAD does not take into account biological localization, presenting simply a fraction of area stained, irrespective of nuclear or cytoplasmic specificity of that particular stain. Cellular segmentation provides a more accurate means of gauging the positivity of cell-associated immunostained biomarkers, as it uses biologically relevant “cell” objects rather than pixels, and intensity-based thresholding on segmented cell objects is a common practice. We tested the ability of MAD and CC, with thresholds to determine low/medium/high intensity staining, for differentiating hypoxic regions within tissue (**Figure 1**). Setting thresholds is justifiable when the marker in question exhibits a binary staining pattern, such as EdU staining in the nucleus, where the presence of any amount of staining would indicate that cell is undergoing DNA synthesis during proliferation. For hypoxia however, the positive hypoxic percentage is greatly influenced by the choice of threshold. The number of cells detected as positive varies from 20-40% of the total cells detected in the viable tissue (**Figure 1D)**, creating a challenge in choosing an appropriate threshold for robust analysis. Calculation of the cumulative histogram of cellular intensity within the image (similar to picking a large range of intensity bins / thresholds) could theoretically be used to model the relationship between a change in intensity and resulting change in area of cells / tissue observed at that intensity. Cumulative histograms for these images are highly nonlinear (also observed in (*Russell et al., 2009*)), leading to difficulty in accurately modeling this relationship (data not shown). Should one wish to perform intensity-based thresholding for markers that exist as gradients within tissue, a judicious choice of thresholding method, or the use of multiple thresholds, along with justification for the particular choice, is warranted, to avoid bias. Vessel distance analysis measures hypoxic marker intensity relative to a known biologically important entity in the tissue, perfused vessel distance, capturing changes in intensity rather than percentage of cells positive at a particular threshold.

While the EF5 positive area may contain regions of both low and high hypoxia, and as such is not the best representation of a spatial- and intensity-varying signal, a ROI-based approach containing a “hypoxic area” can help to define distance relationships between these regions on the slide. Evaluating the area fraction occupied by each ROI (viable tumor, hypoxia and necrosis) versus vessel distance (Figure 2), we obtain an estimate of the width of the hypoxic region, useful either to compare between different treatment strategies, or to evaluate the intrinsic hypoxia tolerance / sensitivity of tumor models. However, a segmentation strategy that looks at individual EF5 cell intensity versus distance would be more accurate than EF5 “region of interest”, due to the intrinsic challenge in either thresholding or training a classifier to detect EF5 “positive” areas.

A common alternative to binary thresholding is using histograms or scatterplots to display the intensity distribution of image pixels. Histograms show distributions of intensities for a marker, grouped either on a per-pixel or per-cell basis. Scatterplot visualization is often used in flow cytometry, where dissociated fluorescently-labelled cells are analyzed for stained intensities of markers of interest. Since this method does not rely on binary thresholds, it provides a useful tool to assess hypoxic gradients. Previous work has shown a strong negative correlation between hypoxia and proliferation using flow cytometry *(Durand and Raleigh, 1998)*. However, flow cytometry lacks the ability to take into account spatial relationships, which are lost upon dissociation of the tissue to single cells. “Tissue cytometry” involves the segmentation and visualization of both intensity and spatial relationships in the cells within an image. **Figure 3** displays two examples of density scatterplot visualizations for EdU and EF5 proliferation and hypoxia, on a per-cell basis. Graphing on a logarithmic scale allows intensities close to the axis to be observed better, and density plots allow for identification of populations of interest. Here, the flow cytometry-like scatterplot of EF5 and EdU marker intensity shows distinct populations of cells that are either hypoxic or proliferating, but not both. As many cyclooxygenases such as COX2 play a role in cellular proliferation *(Sobolewski et al., 2010)*, the hypoxic cells observed were not proliferating. A clear “viable tumor” population exists that is clearly negative for both hypoxia and proliferative markers in the density plot; this could be utilized for objective discrimination between “negative” and “positive” markers. One potential method to quantitatively compare changes in hypoxia and proliferation across different experimental samples would be to gate the population of cells above statistically-determined thresholds (shown as red lines in **Figure 3**) and test if the mean distance of the EF5 or EdU positive cells is significantly different from control groups. If EF5 positive cells display average shorter distances to blood vessels, this could indicate that cells more proximal to perfused vessels may have greater oxygen consumption rates (*Zannella et al., 2013*). If EdU positive cells have shorter distances to blood vessels, this could indicate that higher oxygen concentrations are needed to allow cells to divide and replicate, since proliferating cells use aerobic glycolysis not only for energy but to synthesize intermediates for biosynthetic pathways (*DeBerardinis et al., 2008*). Additionally, coloring each cell by its measured vessel distance (**Figure 3B**) can help to visualize the relationship between distance and hypoxia, at least at a global level. This method, while perhaps difficult to quantify, provides a useful method to interrogate per-cell marker relationships, and may help identify whether markers are correlated or anti-correlated with hypoxia.

To better account for the presence of hypoxic gradients within tissue, we combine the use of distance bins with a cell segmentation methodology for clearer identification of distance relationships. Vessel distance analysis of a KP4 xenograft exemplifies how such data can be presented (**Figure 4)**. We present multiple analyses involving distance to both perfused and all blood vessels, and distance from necrosis, which displays an inverse relationship to the blood vessel distance as expected. While distance to perfused vessel regions is useful to assess from a physiological standpoint, providing a more accurate assessment of chronically hypoxic tumor regions; the use of perfusion dyes is generally only possible in a preclinical setting. However, detection of either total vessel density with CD31 staining, or detection of necrosis, morphologically from H&E tissues, are both possible in clinical specimens. The trends observed in Figure 4, of low EF5 and high EdU staining closer to vessels, and the inverse (high EF5 and low EdU staining closer to necrosis), still hold, indicating that these are viable alternative strategies for hypoxia assessment.

One purpose of distinguishing a perfused vessel subpopulation from all vessels would be to differentiate between chronic and acute hypoxia. Both chronic and acute hypoxia are present in over 50% of solid tumors, but have different clinical implications. For example, acute hypoxia is a greater contributor to genomic instability as opposed to chronic hypoxia. This may be attributed to the generation of reactive oxygen species during periods of reoxygenation of acutely hypoxic regions. In-vivo observations have shown that cells incubated under chronic hypoxia conditions are more invasive than those incubated under acute hypoxia *(Bayer and Vaupel, 2012)*. CD31-positive vessels within the Hoechst ROI (i.e. also positive for Hoechst staining) are considered perfused. Hypoxic gradients relative to perfused vessels would primarily be indicative of chronic hypoxia, due to the balance between diffusion and consumption of oxygen as it exits perfused vessels into surrounding tissue. Hypoxia gradients from perfused vessels measure chronic hypoxia, whereas distance to all vessels measures both chronic and acute hypoxia. Non-perfused vessels occur due to transient vessel occlusion within tumors, leading to the presence of acute hypoxia around these vessels. Each distance analysis provide unique insights about the tumor microenvironment. By combining these analyses, it may even be possible to measure changes in acute hypoxia (i.e. distance to non-perfused/collapsed vessels). However, if there is no need to differentiate the type of hypoxia, distance to all blood vessels is sufficient to capture the hypoxic heterogeneity within the tumor.

To compare distance gradients across multiple samples, one way would be to measure the change in marker intensity across the observed distance of the gradient. In a study, this distance should be constant across both control and experimental (i.e. 200 μm). Calculating the difference in intensity would also correct for background signal, such as EF5 intensity at 10 μm or EdU intensity at 200 μm from a blood vessel. Furthermore, the slopes of the distance gradient in either of the graphs provide different insights. Metabolic oxygen consumption rates could be measured by calculating the slope (calculated as the difference in EF5 intensity divided by distance from perfused or all vessels) in vessel distance analysis; while the cell-intrinsic hypoxia tolerance/sensitivity could be measured by calculating the slope (difference in EF5 intensity divided by distance from necrosis) in necrosis distance analysis. Statistically significant differences in either slope would be indicative of meaningful biological changes, such as a change in oxygen consumption rate or oxygen concentrations needed for cellular division (*Zannella et al., 2013; DeBerardinis et al., 2008*). An alternate method of comparing distance gradients would be to fit the observed curve using a predictive mathematical model, and compare the curve fit parameters across control and experimental groups. Regardless of choice of model used, the change in marker intensity serves biologically meaningful conclusions, providing valuable insight into both oxygen consumption rate and hypoxia tolerance.

Another consideration for clinical immunostaining is the challenge with multiplexing markers on the same tissue slide. The use of serial section immunostaining and alignment / registration of these sections can help compare multiple markers in this setting, though care should be taken with interpretation, due to the presence of different cells in subsequent tissue sections. We simulated this by aligning serial immunofluorescence sections (data not shown). Since hypoxia is present more in regions of low oxygen within the tissue than in particular cells, the proportion of hypoxic staining observed when aligning the DAPI signal from a serial section was similar. However, the number of EdU positive cells observed was greatly reduced, as expected due to the precise intranuclear localization of that marker in proliferating cells. Thus comparing specific colocalization of cell-specific markers would not be recommended, but comparing micro-regional differences in hypoxia, or the proportion of particular cell types on a regional basis, may be possible.

## Conclusion

We have presented several distinct but overlapping methods for analyzing hypoxia and proliferation in solid tumor microenvironments. Each methodology can provide complementary information on the nature of hypoxia within tumors, with different approaches potentially necessary based on the accessibility of markers, and the nature of the scientific question posed. Classification strategies, identifying thresholds for positivity of either pixels or cells, are useful for obtaining an estimate of the percentage of hypoxia within tissues, but suffer from the need to set a specific threshold, which is challenging in the case of a spatially varying signal such as hypoxic gradients. ROI-based distance analyses can be performed on histological images with limited markers, but relies on differences in either tissue morphology or marker intensity to segment these regions. This results in, for example, hypoxic regions of interest containing a range of intensities of the hypoxia marker. Flow cytometry-like scatter plots are useful for visualization and gating of single- and double-negative or positive cell populations, and can be colored by distance to vessel. By segmenting the tissue regions to identify perfused vessels, viable tissue, and necrosis, and calculating per-cell distances to these regions, a distance versus intensity plot can be used to observe changes in cellular phenotypes as a result of decreasing oxygen supply, in order to quantify hypoxia gradients. These methods can be useful to analyze changes in the tumor microenvironment as a result of therapy and as a tool to assess patient hypoxic status in tissue biopsies.

## Contributions to the field / to Associate Editor

Histopathology serves as the gold standard for analysis of hypoxia at the cellular level within tumors. However, though a number of methods are available for measuring hypoxia in tissue sections, many have caveats with analysis or interpretation of the results that can make an accurate assessment of hypoxia challenging. This paper presents a thorough assessment of a number of different methods for visualization and quantification of tumor hypoxia. It will serve as a useful resource for any researchers wishing to perform such studies in the future, by indicating the relative benefits of each visualization / quantification method.

## Author Contributions

MZ, DC, TM, BW contributed to the conception, design, and execution of this study. DC, BW designed and executed the experiments described in this study. MZ, FF, TM developed the analysis methodology with input from all authors. MZ, FF, TM drafted the manuscript; with revisions contributed by all authors. All authors contributed to manuscript revision, read and approved the submitted version.

## Funding

This work was funded by a Terry Fox New Frontiers Research Program (PPG14-1036) grant to BGW, and by an Ontario Graduate Scholarship to DC. The authors would like to acknowledge the Spatio-Temporal Targeting and Amplification of Radiation Response (STTARR) program and its affiliated funding agencies.

## Acknowledgments

The authors would like to thank Dr. Amy Liu and Melania Pintille for helpful discussions and advice on statistical analysis methods.

## Data Availability Statement

The datasets analyzed for this study can be found in the Github repository “Vessel-Distance-Analysis” https://github.com/STTARR/Vessel-Distance-Analysis

## Notes

https://github.com/STTARR/Vessel-Distance-Analysis

## References

1. Alper, T., & Bryant, P. E. (1974). Reduction in oxygen enhancement ratio with increase in LET: Tests of two hypotheses. International Journal of Radiation Biology and Related Studies in Physics, Chemistry and Medicine, 26(3), 203–218.

2. Gomez, C. R. (2016). tumor Hypoxia: impact in tumorigenesis, diagnosis, Prognosis, and therapeutics. Frontiers in oncology, 6, 229.

3. Milosevic, M. F., Fyles, A. W., & Hill, R. P. (1999). The relationship between elevated interstitial fluid pressure and blood flow in tumors: a bioengineering analysis. International Journal of Radiation Oncology* Biology* Physics, 43(5), 1111–1123.

4. Thomlinson, R. H., & Gray, L. H. (1955). The histological structure of some human lung cancers and the possible implications for radiotherapy. British journal of cancer, 9(4), 539.

5. Haugland, H. K., Vukovic, V., Pintilie, M., Fyles, A. W., Milosevic, M., Hill, R. P., & Hedley, D. W. (2002). Expression of hypoxia-inducible factor-1α in cervical carcinomas: correlation with tumor oxygenation. International Journal of Radiation Oncology* Biology* Physics, 53(4), 854–861.

6. Bayer, C., & Vaupel, P. (2012). Acute versus chronic hypoxia in tumors. Strahlentherapie und Onkologie, 188(7), 616–627.

7. Vaupel, P., & Mayer, A. (2007). Hypoxia in cancer: significance and impact on clinical outcome. Cancer and Metastasis Reviews, 26(2), 225–239.

8. Jensen, R. L. (2009). Brain tumor hypoxia: tumorigenesis, angiogenesis, imaging, pseudoprogression, and as a therapeutic target. Journal of neuro-oncology, 92(3), 317–335.

9. Carreau, A., Hafnyl◻Rahbi, B. E., Matejuk, A., Grillon, C., & Kieda, C. (2011). Why is the partial oxygen pressure of human tissues a crucial parameter? Small molecules and hypoxia. Journal of cellular and molecular medicine, 15(6), 1239–1253.

10. Rudat, V., Stadler, P., Becker, A., Vanselow, B., Dietz, A., Wannenmacher, M., … & Feldmann, H. J. (2001). Predictive value of the tumor oxygenation by means of pO2 histography in patients with advanced head and neck cancer. Strahlentherapie und Onkologie, 177(9), 462–468.

11. Milosevic, M., Warde, P., Ménard, C., Chung, P., Toi, A., Ishkanian, A., … & Bristow, R. (2012). Tumor hypoxia predicts biochemical failure following radiotherapy for clinically localized prostate cancer. Clin Cancer Res, 18(7), 2018–14. doi: 10.1158/1078-0432.CCR-11-2711

12. Mirabello, V., Cortezon-Tamarit, F., & Pascu, S. I. (2018). Oxygen Sensing, Hypoxia Tracing and in Vivo Imaging with Functional Metalloprobes for the Early Detection of Non-communicable Diseases. Frontiers in chemistry, 6, 27.

13. Loukas, C. G., & Linney, A. (2004). A survey on histological image analysis-based assessment of three major biological factors influencing radiotherapy: proliferation, hypoxia and vasculature. Computer Methods and Programs in Biomedicine, 74(3), 183–199.

14. Russell, J., Carlin, S., Burke, S. A., Wen, B., Yang, K. M., & Ling, C. C. (2009). Immunohistochemical detection of changes in tumor hypoxia. International Journal of Radiation Oncology* Biology* Physics, 73(4), 1177–1186.

15. Urtasun, R. C., Chapman, J. D., Raleigh, J. A., Franko, A. J., & Koch, C. T. (1986). Binding of 3H-misonidazole to solid human tumors as a measure of tumor hypoxia. International Journal of Radiation Oncology* Biology* Physics, 12(7), 1263–1267.

16. Rijken, P. F., Bernsen, H. J., Peters, J. P., Hodgkiss, R. J., Raleigh, J. A., & van der Kogel, A. J. (2000). Spatial relationship between hypoxia and the (perfused) vascular network in a human glioma xenograft: a quantitative multi-parameter analysis. International Journal of Radiation Oncology* Biology* Physics, 48(2), 571–582.

17. Wijffels, K. I. E. M., Kaanders, J. H. A. M., Rijken, P. F. J. W., Bussink, J., Van Den Hoogen, F. J. A., Marres, H. A. M., … & Van Der Kogel, A. J. (2000). Vascular architecture and hypoxic profiles in human head and neck squamous cell carcinomas. British journal of cancer, 83(5), 674.

18. Swinson, Daniel EB, J. Louise Jones, Donna Richardson, Charles Wykoff, Helen Turley, Jaromir Pastorek, Nick Taub, Adrian L. Harris, and Kenneth J. O’Byrne. “Carbonic anhydrase IX expression, a novel surrogate marker of tumor hypoxia, is associated with a poor prognosis in non–small-cell lung cancer.” Journal of Clinical Oncology 21, no. 3 (2003): 473–482.

19. Primeau, A. J., Rendon, A., Hedley, D., Lilge, L., & Tannock, I. F. (2005). The distribution of the anticancer drug Doxorubicin in relation to blood vessels in solid tumors. Clinical Cancer Research, 11(24), 8782–8788.

20. Tannock, I. F. (1968). The relation between cell proliferation and the vascular system in a transplanted mouse mammary tumour. British journal of cancer, 22(2), 258.

21. Cojocari, D. (2017). Therapeutic Targeting of Tumour Hypoxia through the Unfolded Protein Response and Autophagy Pathways (Doctoral dissertation, University of Toronto).

22. Durand, R. E., & Raleigh, J. A. (1998). Identification of nonproliferating but viable hypoxic tumor cells in vivo. Cancer research, 58(16), 3547–3550.

23. Sobolewski, C., Cerella, C., Dicato, M., Ghibelli, L., & Diederich, M. (2010). The role of cyclooxygenase-2 in cell proliferation and cell death in human malignancies. International journal of cell biology, 2010.

24. Zannella, V. E., Dal Pra, A., Muaddi, H., McKee, T. D., Stapleton, S., Sykes, J., … & Wouters, B. G. (2013). Reprogramming metabolism with metformin improves tumor oxygenation and radiotherapy response. Clinical cancer research, 19(24), 6741–6750.

25. DeBerardinis, R. J., Lum, J. J., Hatzivassiliou, G., & Thompson, C. B. (2008). The biology of cancer: metabolic reprogramming fuels cell growth and proliferation. Cell metabolism, 7(1), 11–20.

26. Haynes J., McKee T. D., Haller A., Wang Y., Leung C., Gendoo D. M. A., … & O’Brien C. A. (2018). Administration of Hypoxia-Activated Prodrug Evofosfamide after Conventional Adjuvant Therapy Enhances Therapeutic Outcome and Targets Cancer-Initiating Cells in Preclinical Models of Colorectal Cancer. Clinical Cancer Research, 24(9):2116–2127; DOI:10.1158/1078-0432.CCR-17-1715

