## Supplemental Figure 1 for "Quantitative Visualization of Hypoxia and Proliferation Gradients Within Histological Tissue Sections"

Supplementary Material

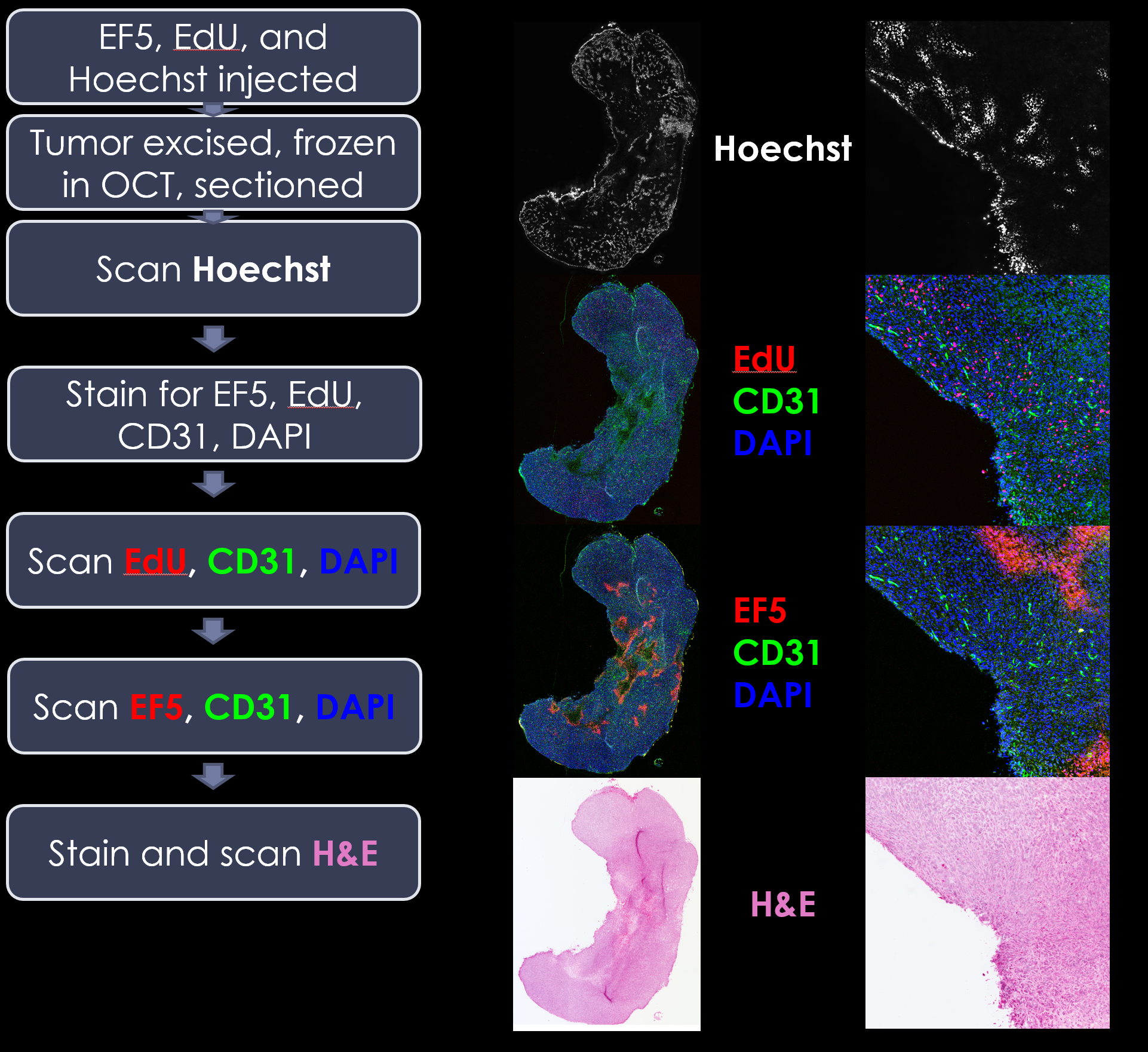

***Supplementary Figure 1.*** ***Immunofluorescent histology and imaging workflow.*** *Flowchart (left) depicting process of alternating scanning and staining needed to acquire the four images (right) utilized for subsequent image processing and analysis of perfused vessel distance, proliferation and hypoxia.*
